# PhylloTraits 1.0: Unveiling the diversity of functional traits

**DOI:** 10.1101/2025.05.13.653771

**Authors:** Federico Villalobos, Romeo A. Saldaña-Vázquez, Yara Azofeifa-Romero, Willy Pineda-Lizano, Mariano S. Sánchez, Jorge Ayala-Berdon, Jorge D. Carballo-Morales, John Harold Castaño, Jesús R. Hernández-Montero, Leonel Herrera-Alsina

## Abstract

Functional traits play a key role in understanding species’ ecological and evolutionary dynamics. However, the plenty of data collected on functional traits is sparse across literature so retrieving it, especially information for tropical species, becomes a challenge. We introduce PhylloTraits, a functional trait comprehensive database from bats of the family Phyllostomidae. The New World phyllostomids are one of the mammalian families with the greatest diversity of trophic and ecological habits. In addition, this family of bats is a prominent component of mammalian assemblages in the Neotropics. Phyllotraits 1.0 compiles a data collection on wing and body morphology from 230 species at individual level. The wing morphology traits can provide insight into flight performance and maneuverability. The body morphology traits are useful in addressing ecological aspects of Phyllostomidae bats. PhylloTraits 1.0 provides efficient access to functional data collated from various sources, including published literature in English, Spanish and Portuguese, and field studies. Furthermore, PhylloTraits 1.0 includes geographic coordinates of individual data to facilitate the examination of biogeographic patterns across different regions. By uncovering and synthesizing these functional traits and capturing individual variation, PhylloTraits aims to enhance our understanding of the ecological roles, evolutionary adaptations, and conservation implications of phyllostomid bats. This database would serve as a tool for comparative studies, ecological modeling, and conservation assessments. We anticipate that PhylloTraits will stimulate further research and promote collaborative efforts to unravel the intricate relationships between functional traits, their evolution and the ecological dynamics of Phyllostomidae bats.

## Introduction

Functional traits include physiological, anatomical, biochemical, phenotypic, and behavioral characteristics of organisms directly related to their function in ecosystems (Violle et al., 2007). These traits are crucial for understanding the ecology and evolutionary biology of species as they influence their survival, reproduction, and adaptation to the environment (Díaz et al., 2013). The importance of functional traits in ecological and evolutionary biology studies lies in their ability to explain how organisms interact with their surroundings, adapt to different environmental conditions, and evolve over time (Díaz et al., 2013).

Functional trait databases enable researchers to systematically access a wealth of information, facilitating comparisons between species and identifying patterns and trends (Froidevaux et al., 2023). Importantly, some existing databases lack geographic information, limiting their use to analyze both ecological and biogeographic patterns (Table 1). For instance, by incorporating geographic information into functional trait databases, we are able to study ecological interactions in a spatially-explicit context to study the historical patterns that have shaped the morphological diversity of this group and to correlate variation in functional traits to environmental variation (Thompson, 2005).

**Table 1.**
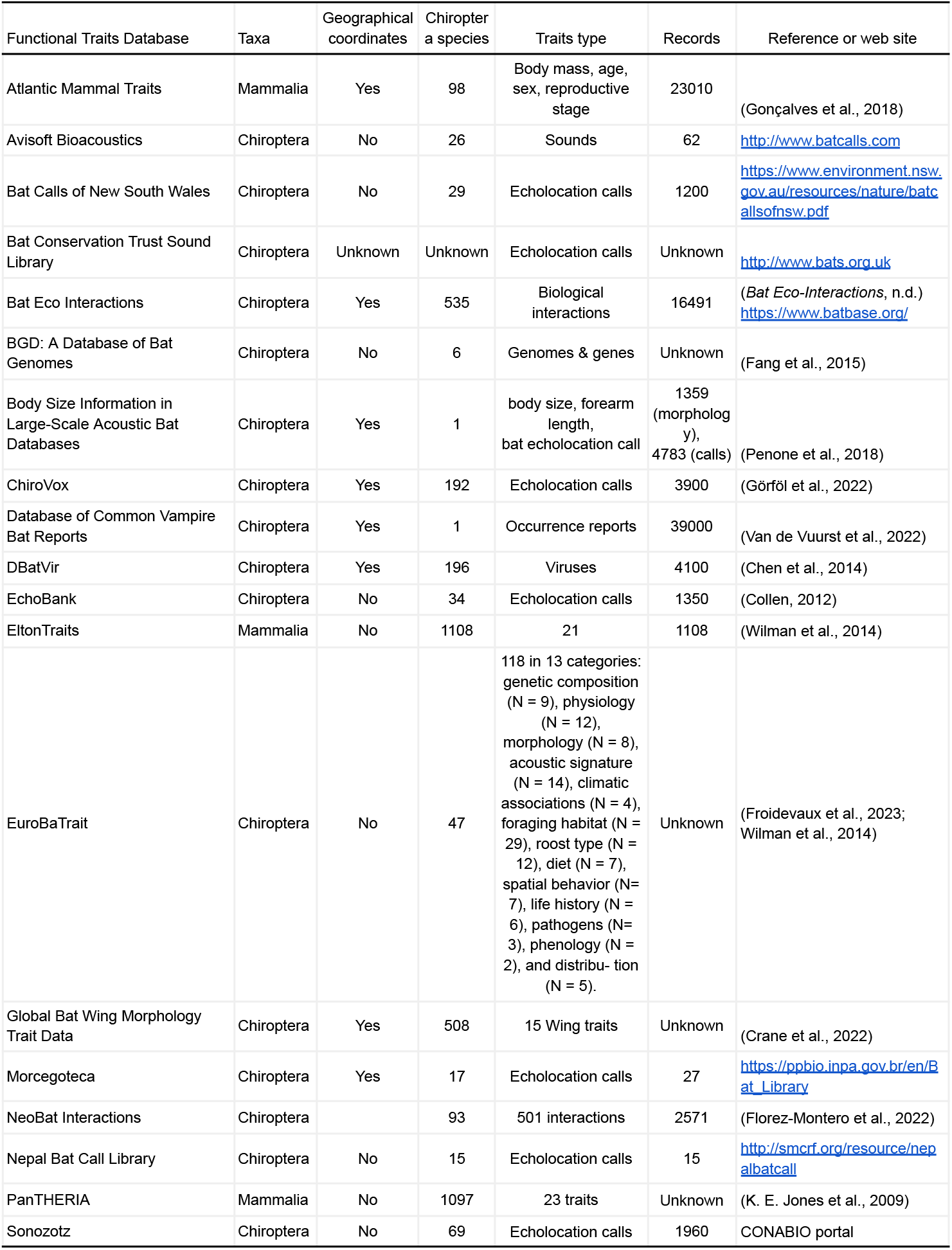
Features of public available Chiroptera databases

Bats exhibit unique adaptations such as membranous wings, sophisticated echolocation systems, and a wide diversity of feeding types, allowing them to occupy a variety of ecological niches. For these flying mammals, the study of functional traits provides insights into characteristics associated with flight capacity, feeding habits, echolocation, and reproduction. Within the order Chiroptera, the Phyllostomidae family is highly diverse in species representing different ecological niches. Restricted to the tropical and subtropical regions of the Americas, a remarkable evolutionary radiation resulted in 230 species and 60 genera with an exceptional diversity of ecological and trophic habits that include frugivorous, insectivorous, nectarivorous, carnivorous, and hematophagous species (Fleming et al., 2020). These feeding types are associated with a broad variation in morphological traits, for example, body mass ranges from 4.72 g to 167.5 g (Moyers Arévalo et al., 2020). Phyllostomid bats play a crucial role in Neotropical ecosystems, serving as important plant pollinators, insect population regulators and seed dispersers (Kunz et al., 2011). Their diversity and adaptability make them a fascinating group for research in ecology, evolution, and conservation. Finally, Phyllostomid bats are also considered bioindicators of ecosystem health, as their presence and diversity reflect environmental quality and stability (Medellín et al., 2000; G. Jones et al., 2009).

In addition to their ecological importance, the Phyllostomidae family is among the most extensively studied bat groups in the Neotropics from an ecological perspective (Fig. 1). This is mostly due to their dominance in Neotropical bat communities and the relative high probability of capturing them with mist nets. However, the available knowledge on this family is fragmented, scattered across various sources, including scientific articles, undergraduate theses, grey literature, and unpublished field data. The aim of this paper is to introduce the Phyllostomid bat functional trait database, ‘PhyloTraits 1.0,’ which consolidates existing information and identifies knowledge gaps regarding phyllostomid species. We expect this database to facilitate the analysis of biogeographic and ecological patterns, as well as comparative biology within the Phyllostomidae family

**Figure 1.**
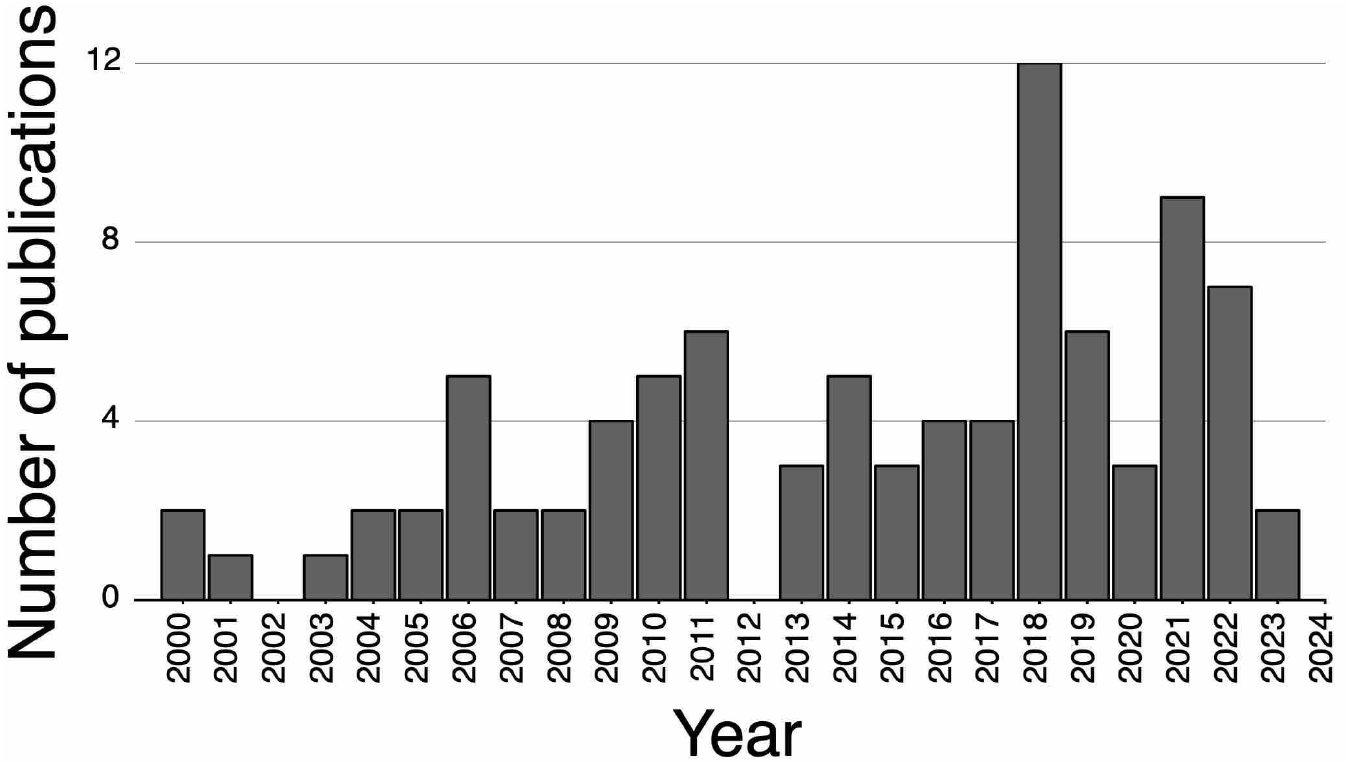
Publications per year for Phyllostomidae in the area of ecology, according to WoS (2023).

## Materials and methods

Functional traits in bats are divided into two major groups: morphological and life history traits (Castillo-Figueroa & Pérez-Torres, 2021). Morphological traits are linked to the characteristics that enable bats to interact differently with the environment, associated with flight speed, foraging habitat, and foraging strategy. Life history traits are related to reproduction, physiology, behavior, and the dimensions of the ecological niche they occupy.

For this initial version of PhylloTraits, we focused on 17 morphological traits (Table 2). These traits were collected from scientific literature which also include published papers in local journals, gray literature and undergrad thesis. Most of these items were originally published in Spanish which complicates its accessibility for international researchers. In future versions the database will include information from 1) field measurements, 2) biological collections, and 3) data obtained from biophysical models.

**Table 2.** Functional traits computed by PhylloTraits 1.0.

| Trait type | Functional dimension | Trait |
| --- | --- | --- |
| Morphologic | Size | body mass, total length. |
|  | Wings | tip shape index, wing span, wing loading, length of the third and fifth digit. |
|  | Head | length of nose leaf, length of the ear, length of tragus. |
|  | Hindlimbs | length of the foot, length of the tibia. |
|  | Pollex/thumb | length of the pollex/thumb |

To collate information from published scientific studies, we conducted 17 systematic literature searches on Google Scholar. The keywords, search engine, and the number of results for each search are described in Supplementary Table 1. Additionally, secondary literature searches were conducted based on studies collected in the systematic review. The searches were performed in Portuguese, Spanish, and English. Data entry was performed at an individual level whenever this information was available for each trait. In addition to individual trait values, supplementary information was included, such as the nature of data source (e.g., bibliographic, collections, field), statistics (means, SD, sample sizes, and range), geographic information (e.g., locality, province/state, country, coordinates) and sex of the individuals when available.

With the information gathered, a series of completeness statistics by species, genus, and distribution area are presented below. The geographic range size of species was obtained from Rojas et al.(2018).

## Results

The DataBase is composed of 254 species and 2445 individual records from 141 published scientific reports, and 159 records from unpublished field data, and 37 records from natural history collections. Regarding the information richness for each trait, the most extensively documented trait was foot length, followed by traits commonly used for species identification in the field, such as forearm length, body mass, total length, and ear length. In contrast, data on wing surface and the length of the fifth digit were each derived from a single reference.

The degree of data completeness is variable between traits. We found that traits associated with the identification of species (body mass, ear length and total length) were easier to collate than traits that describe wing morphology (e.g, length of each phalange) (Fig. 2)

**Figure 2.**
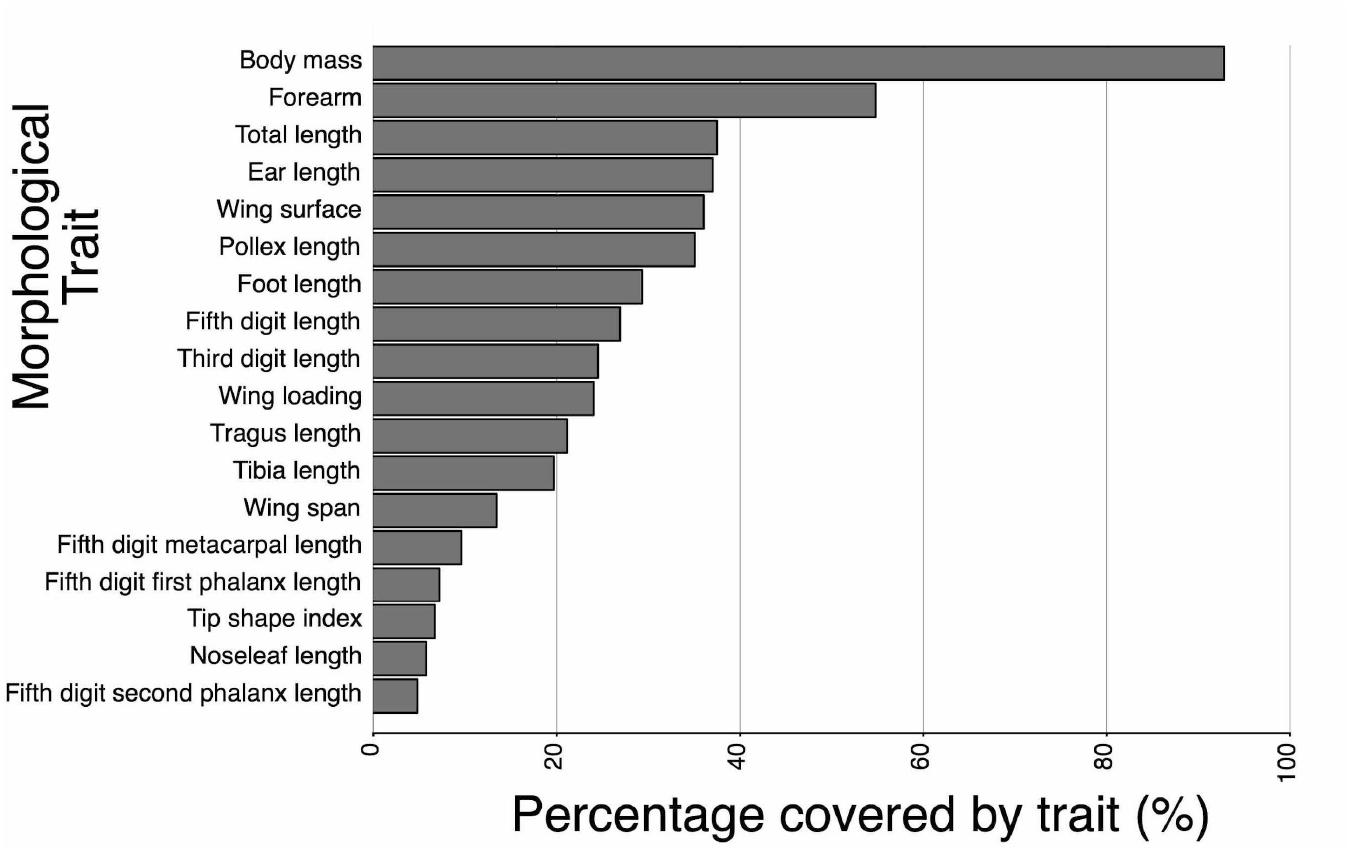
Percentage of trait completeness

We found that only 6% of the Phyllostomid species have > 70% of trait completeness. Prominently within this category are species such as *Artibeus jamaicensis, Carollia perspicillata*, and *Carollia castanea*. For the 36% of species, less than 10% of functional traits are known (Fig. 3)..

**Figure 3.**
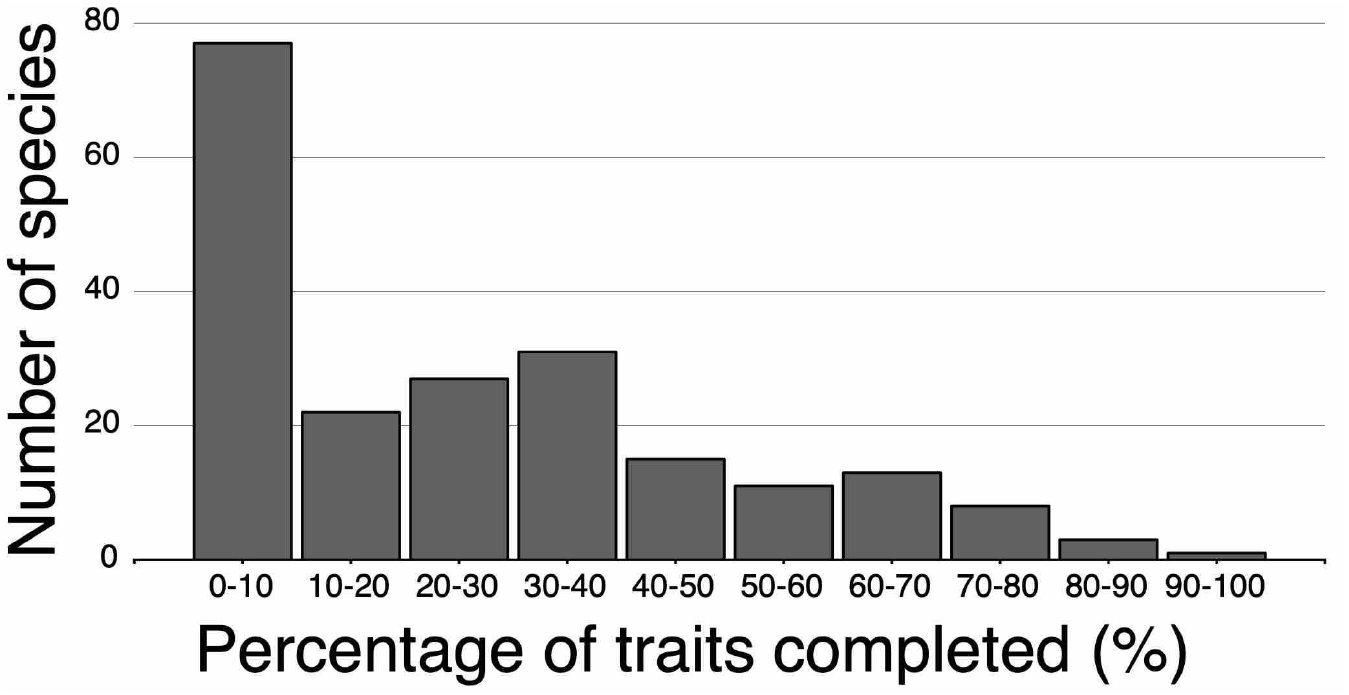
Traits percentage available by species number.

For 71% of genera, we found less than 70% of data for all traits (see Fig. 4). A specific set of 12 (20%) genera are characterized by exceptionally low information levels, falling below15%. Notably, the genus *Musonycteris*, and *Monophyllus*, endemic to Mexico and Antilles, are particularly lacking trait data.

**Figure 4.**
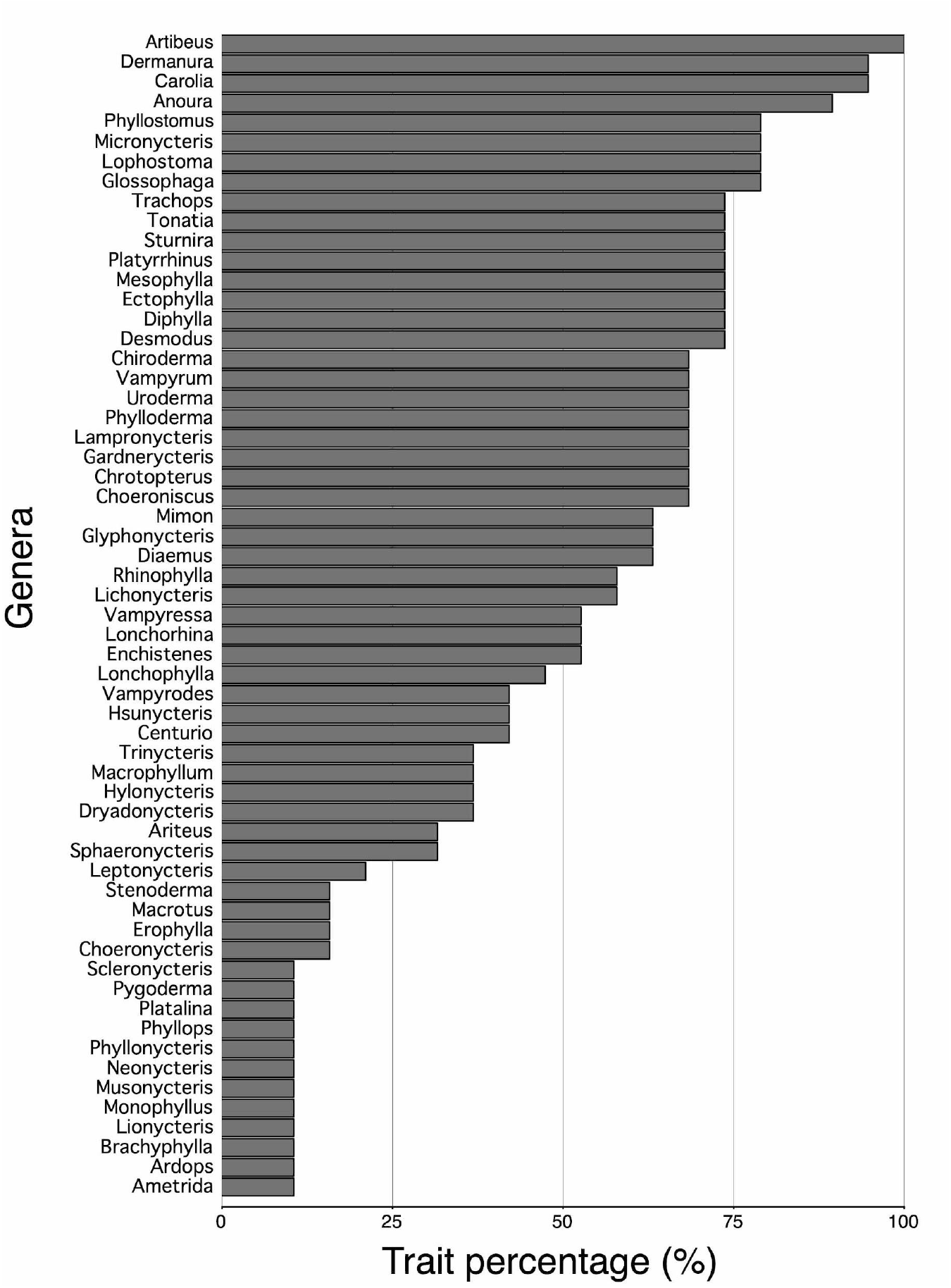
Completeness percentage trait per genus.

We found that species with larger geographic ranges tend to have more available trait information in literature (Figure 4). However, the relationship between these two variables was relatively weak (*R*^2^ = 0.37; see Fig. 5). Species with restricted distribution show considerable information levels, such as *Artibeus jamaicensis* and *Carollia castanea*. In contrast, the nectarivorous bat *Lionycteris spurrelli*, despite its wide distribution range, has an information completeness of less than 0.5%.

**Figure 5.**
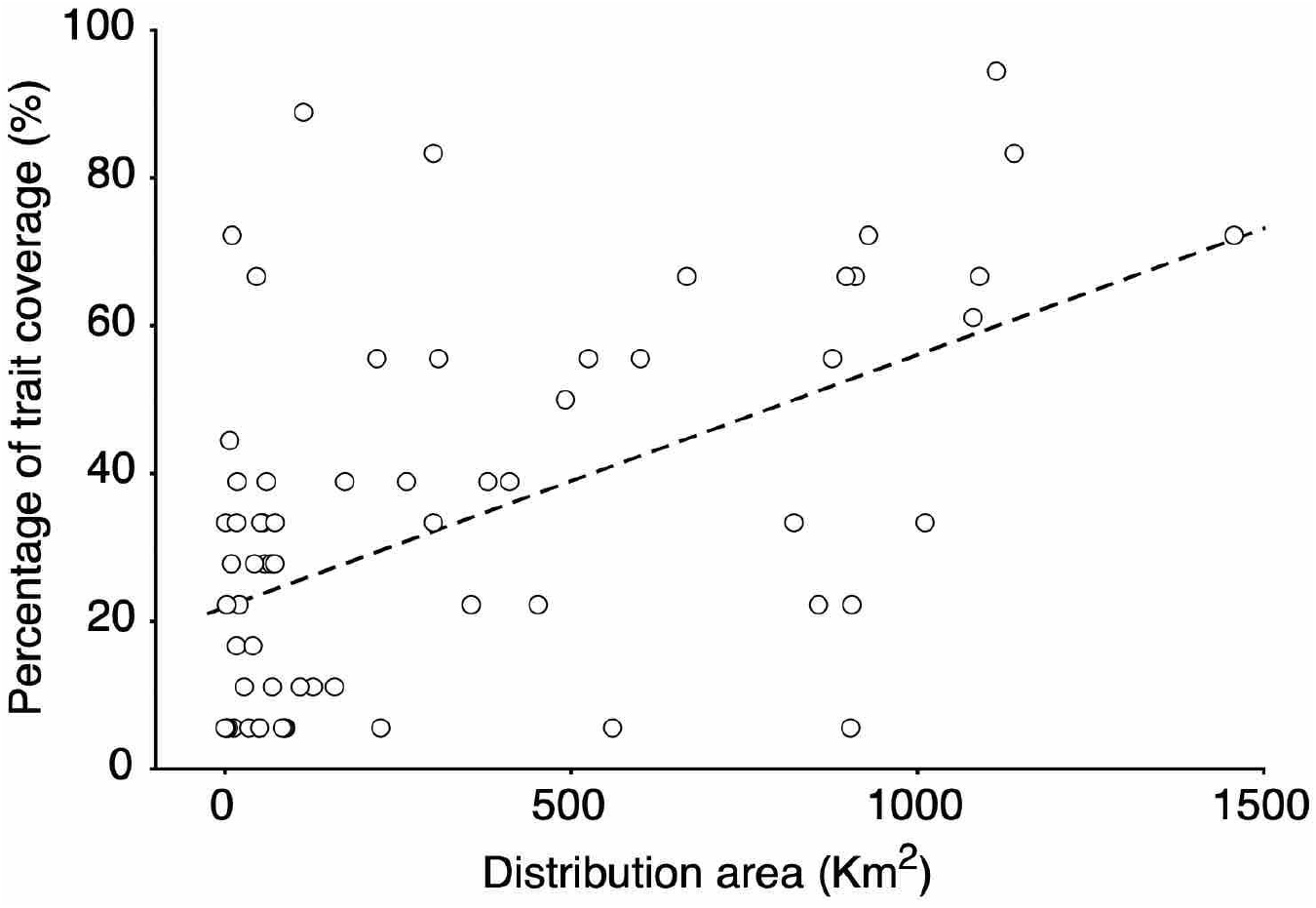
Relationship between distribution size of phyllostomid species and the percentage of traits coverage.

## Discussion

PhylloTraits 1.0 offers a valuable contribution to the field of bat ecology and evolution by providing a comprehensive resource for exploring the functional traits of Phyllostomidae bats. Here, we discuss the significance of this database, address potential explanations for information gaps, and explore future directions for enriching and utilizing this resource.

### Significance of PhylloTraits in Bat Research

PhylloTraits 1.0 stands out from existing databases in several key ways. First, it is focused on a specific and ecologically important bat family which provides a deeper and more detailed understanding of the functional traits within this diverse group. Second, because PhylloTraits focuses on a monophyletic group, analyses on speciation and extinction rates are statistically more robust than analyses done on non-monophyletic groups. Third,, the future integration of data from multiple sources (i.e., published literature, museum collections, field studies) will offer a complete picture of the ecological and macroevolutionary patterns. Fourth, by integrating functional and geographic information, our database enables the examination of biogeographic patterns and ecological interactions across different regions.

PhylloTraits 1.0 provides the opportunity to tackle a range of ecological and evolutionary questions related to phyllostomid bats, such as: understanding how functional traits influence niche occupation and resource partitioning within the family (Chang et al., 2019); (Jiang et al., 2018); investigating the drivers of diversification and adaptation across different ecological contexts (Potter et al., 2021); (Rojas et al., 2012); assessing the vulnerability of Phyllostomidae species to environmental changes and anthropogenic threats (Carballo-Morales et al., 2021); and informing conservation and management strategies for these ecologically relevant bats.

We have identified information gaps at different taxonomic levels. Some species, due to their abundance and wide distribution, have multiple sources of information, whereas others have very limited data available in the literature for the selected traits. This information gap is also evident at the level of the functional traits evaluated. Traits associated with taxonomic identification are better documented, while more specialized traits (e.g., aerodynamic characteristics) are less well studied. By integrating data on key functional traits, this database addresses a critical information gap in bat research, particularly in the Neotropics. The comprehensive nature of PhylloTraits, encompassing body and wing morphology, offers an integrated view of the functional ecology of these bats as suggested by (Castillo-Figueroa & Pérez-Torres, 2021). This is particularly crucial given the role of Phyllostomidae bats in ecosystem functions such as pollination and seed dispersal (González-Gutiérrez et al., 2022).

### Insights into Ecological and Evolutionary Patterns

The database will facilitate a deeper understanding of the ecological roles and evolutionary adaptations of Phyllostomidae bats. For instance, the variation in wing morphology across species can be linked to differences in flight performance and foraging strategies, reflecting ecological niche specialization (García-Herrera et al., 2022); (Ospina-Garcés et al., 2023). By focusing on traits like wing morphology, PhylloTraits provides insights into how these bats interact with their environment and adapt to different ecological niches. In addition, skeletal morphology data can provide insights into flight biomechanics and foraging habits, terrestrial locomotion, or roosting behavior (Louzada et al., 2019; Sánchez & Carrizo, 2021). This information is vital for understanding how these bats have evolved in response to different ecological pressures. Functional traits are not just static measurements but are connected to the ecological roles and evolutionary strategies of species (Violle et al., 2007). For the future and more ambitious version of PhylloTraits, we are already collecting data on skull morphology which will provide insights into feeding behaviors and dietary adaptations (García-Herrera et al., 2021).

### Phylogenetic Context and Ecosystem Services

Understanding how the ecology and evolution of functional traits affect species’ roles in ecosystems and their resilience to environmental changes is essential. The phylogenetic context is critical for assessing ecosystem service vulnerability (Díaz et al., 2013). PhylloTraits can play a key role in exploring the phylogenetic aspects of functional traits in bats (e.g., (Cabrera-Campos et al., 2021). This is crucial for understanding how changes in bat functional traits may impact populations and distributions, and consequently, ecosystem services such as pollination and seed dispersal, which are vital in tropical ecosystems.

### Potential for Comparative Studies and Ecological Modeling

PhylloTraits 1.0 offers potential for broader ecological and evolutionary studies. The database can be used for comparative studies within the Phyllostomidae and across different bat families or other mammalian groups, enhancing our understanding of mammalian ecology and evolution. Furthermore, the data might be integrated into ecological models to predict how changes in environmental conditions might impact these bats, which is essential for anticipating the effects of climate change and habitat alteration (e.g. (Ortega-García et al., 2017)).

### Biogeographic Patterns and Conservation Implications

The inclusion of a spatial dimension in PhylloTraits will allow for the examination of biogeographic patterns. Spatial information is not available in all existing databases focused on Chiroptera (only on 42% of reviewed databases). This aspect is particularly important for understanding how environmental factors and habitat changes influence the distribution and diversity of Phyllostomidae bats. Such insights are crucial for conservation efforts, especially in the face of habitat loss and climate change (Carballo-Morales et al., 2021). Although the distributions of the Phyllostomidae have been modelled (Rojas et al., 2018), the spatial mapping of functional traits for these species has not been systematically compiled. The database can help identify species and regions that are particularly vulnerable, guiding targeted conservation actions.

### Information Gaps and Explanatory Hypotheses

Despite its comprehensiveness, PhylloTraits 1.0 reveals significant information gaps for various species and traits(i.e., *Musonycteris, Monophyllus, Lionycteris)*. Understanding the factors behind these gaps is crucial for improving the database and maximizing its utility. Here, we propose several hypotheses:

1. Species abundance: Rare or less abundant species are inherently less likely to be encountered and studied, leading to gaps in trait data.
2. Reduced geographic distribution: Species with restricted ranges are naturally less accessible for research, potentially contributing to information deficiencies.
3. Trait complexity: Certain traits, particularly those requiring specialized measurements or equipment, may be more challenging to record, leading to data sparsity (i.e. landmarks, wing morphology).
4. Taxonomic uncertainty: Uncertainties in species identification or classification can hinder data collection and integration (i.e. *Sturnira* species names have recently changed, which must be taken into account when retrieving data from older publications containing basic information).
5. Research bias: Geographical and taxonomic biases exist in scientific research, with certain regions and charismatic species receiving more attention than others. This can lead to uneven data distribution within the database. Charismatic and umbrella species (e.g., *Ectophylla alba* Honduran white bat) may attract more attention and be better researched due to their unique features or ecological roles, potentially inflating their information content in the database. Also there may be an institutional bias. Institutions with greater research capacity and resources may contribute more data to the database, potentially skewing representation towards specific regions or taxa. For example, La Selva biological station in Costa Rica and the STRI Smithsonian in Panama might have contributed more data on bat species found in the region compared to less-studied regions.

The distribution of information gaps within PhylloTraits 1.0 is not random. We observe a trend of lower information content for species with restricted geographic ranges (Figure 5), highlighting the need for targeted research efforts in areas with understudied bat diversity. These information gaps may manifest differently at the trait, genus, and species levels. At the trait level, certain traits, particularly those requiring specialized expertise or equipment, may exhibit more significant gaps across the board. We also expect at the species level that rare, geographically restricted, or taxonomically uncertain species are most likely to have substantial information gaps in the database. Also, genera with a high number of rare or geographically restricted species are likely to have lower overall information content. Geographic distribution and institutional biases might be more pronounced at the genus and species levels, where certain groups are studied more intensively due to their distribution around research centers or their ecological significance.

### Perspectives and Recommendations

Our analysis reveals significant gaps in the available data, particularly for certain species and traits. This uneven distribution of information highlights the need for more targeted research efforts, especially for underrepresented species and traits. Future versions of PhylloTraits should aim to fill these gaps, we believe that collaboration and data sharing will be an effective strategy to fill data gaps and ensure wider representation in the database. Additionally, the database could be expanded to include more life history traits, which would provide a more comprehensive understanding of the evolutionary dynamics of these bats.

To address these gaps and biases, future efforts should focus on:

1. Targeted research: Prioritizing underrepresented species and traits, especially in understudied regions, to balance data representation. Institutional biases might be geographically clustered, leading to data deserts in regions with limited research infrastructure or capacity (e.g., Nicaragua in Central America). Addressing these geographical and institutional imbalances is crucial for ensuring a more comprehensive and representative database.
2. Collaborative efforts: Encouraging collaborations between institutions from different regions in a micro and macro scope can help diversify the data sources and reduce geographical biases.
3. Citizen science and remote sensing: Utilizing citizen science and remote sensing technologies can help gather data on hard-to-reach species or regions.
4. Standardization and open access: Promoting standardization in data collection and ensuring open access to data can facilitate broader participation and usage by the global research community.

## Supplementary information

Supplementary Table 1: Keywords and the number of documents results for each search.

PhylloTraits 1.0 database is available at: https://doi.org/10.5281/zenodo.15272507

## Acknowledgments

Romeo A. Saldaña-Vázquez would like to thank the Universidad Iberoamericana Puebla for the facilities provided for the completion of this work and the research stays at the Universidad Nacional de Costa Rica, as well as Federico Villalobos’ research stays at the Universidad Iberoamericana Puebla. Mariano S. Sánchez to thank the Consejo Nacional de Investigaciones Científicas y Técnicas (CONICET) for institutional support and Fondo para la Investigación Científica y Tecnológica (FONCYT) for research grants PICT 2020-03352. Phyllotraits Database is adscribe to Laboratorio de Sistemática, Genetica y Evolucion (LabSGE, project number PPAA 0142-22) at Universidad Nacional de Costa Rica, and funded by Fondo especial para la educación superior de Costa Rica (FESS).

